# A long reads-based *de-novo* assembly of the genome of the Arlee homozygous line reveals structural genome variation in rainbow trout

**DOI:** 10.1101/2020.12.28.424581

**Authors:** Guangtu Gao, Susana Magadan, Geoffrey C. Waldbieser, Ramey C. Youngblood, Paul A. Wheeler, Brian E. Scheffler, Gary H. Thorgaard, Yniv Palti

## Abstract

Currently, there is still a need to improve the contiguity of the rainbow trout reference genome and to use multiple genetic backgrounds that will represent the genetic diversity of this species. The Arlee doubled haploid line was originated from a domesticated hatchery strain that was originally collected from the northern California coast. The Canu pipeline was used to generate the Arlee line genome de-novo assembly from high coverage PacBio long-reads sequence data. The assembly was further improved with Bionano optical maps and Hi-C proximity ligation sequence data to generate 32 major scaffolds corresponding to the karyotype of the Arlee line (2N=64). It is composed of 938 scaffolds with N50 of 39.16 Mb and a total length of 2.33 Gb, of which ∼95% was in 32 chromosome sequences with only 438 gaps between contigs and scaffolds. In rainbow trout the haploid chromosome number can vary from 29 to 32. In the Arlee karyotype the haploid chromosome number is 32 because chromosomes Omy04, 14 and 25 are divided into six acrocentric chromosomes. Additional structural variations that were identified in the Arlee genome included the major inversions on chromosomes Omy05 and Omy20 and additional 15 smaller inversions that will require further validation. This is also the first rainbow trout genome assembly that includes a scaffold with the sex-determination gene (sdY) in the chromosome Y sequence. The utility of this genome assembly is demonstrated through the improved annotation of the duplicated genome loci that harbor the IGH genes on chromosomes Omy12 and Omy13.

**Article Summary:** A *de-novo* genome assembly was generated for the Arlee homozygous line of rainbow trout to enable identification and characterization of genome variants towards developing a rainbow trout pan-genome reference. The new assembly was generated using the PacBio sequencing technology and scaffolding with Hi-C contact maps and Bionano optical mapping. A contiguous genome assembly was obtained, with the contig and scaffold N50 over 15.6 Mb and 39 Mb, respectively, and 95% of the assembly in chromosome sequences. The utility of this genome assembly is demonstrated through the improved annotation of the duplicated genome loci that harbor the IGH genes.

## Introduction

Because of its high economic value for sport fishing and aquaculture industries, rainbow trout (*Oncorhynchus mykiss*) is the most cultivated cold-water fish in the US and one of the most studied aquaculture species. As model research organism, rainbow trout have also been used to study comparative immunology, evolution and development, ecology and to study the carcinogenicity of several food and environmental contaminants. Genomics studies have been important in conducting discovery genotype to phenotype research and to define genetic parameters that control complex traits such as disease resistance and feed efficiency in aquaculture breeding programs. A well-annotated reference genome assembly is a vital element in genome-wide association studies and for transcriptome and proteome expression studies. For rainbow trout, there were two versions of genome assemblies available in NCBI prior to the assembly we present in this report. Both versions were based on short-read sequencing technologies, with the AUL_PRJEB4421_v1 assembly (NCBI accession GCA_900005704.1) (Berthelot et al. 2014) built using 454 pyrosequencing and Illumina data and the Omyk_1.0 assembly (NCBI accession GCA_002163495.1) (Pearse et al. 2019) which was based on Illumina sequencing data.

The genome size estimate for rainbow trout based on cellular DNA content is 2.4 Gb (Hardie and Hebert 2003). It is larger than most economically important fish species and aquatic model research organisms. Although it is similar in size to that of most mammals, its architecture and organization are more complex because it underwent two additional genome duplication events and has higher content of long repeats. All ray-finned fish share an additional (3R) round of ancestral genome duplication in their evolutionary history compared to mammals and birds, but the salmonids underwent an additional (4R) whole genome duplication event (Allendorf and Thorgaard 1984; Lien et al. 2016). In total, ∼90% of the rainbow trout genome is composed of 88 pairs of colinear blocks of high sequence similarity known as homeologous chromosome regions (Allendorf et al. 2015), including seven pairs of chromosome arms with elevated sequence similarity (>90%) and patterns of tetrasomic inheritance (Pearse et al. 2019). In addition, ∼57% of the rainbow trout genome contains repetitive sequences (Pearse et al. 2019; Genet et al. 2011). This high level of repetitive sequences reduces the contiguity of de-novo genome assemblies that are based on short-read sequencing data leading to high number of contigs with many ambiguous gaps between the assembled contigs. Moreover, many genome segments that contain long repetitive sequences can collapse into few assembly contigs due to their high similarity and the inability of the short reads to span the long repeat sequences. For the current rainbow trout genome assemblies, there are 141,187 spanned and 120 unspanned gaps in the AUL_PRJEB4421_v1 assembly and 420,055 spanned and 7,839 unspanned gaps in the Omyk_1.0 assembly. The assembly gaps and the mis-assembled repetitive sequences also make it challenging to scaffold the assembly contigs and then to anchor the scaffolds to the chromosomes. In the AUL_PRJEB4421_v1 assembly, the scaffold N50 is only 383,627bp and ∼95% of the assembly sequence is not mapped to chromosome sequences. Although a significant improvement over the AUL_PRJEB4421_v1 assembly, the Omyk_1.0 assembly’s N50 is still modest (1,670,138bp) and over 10% of the assembly bases are in 131,931 unplaced scaffolds. By spanning the repetitive regions and resolving duplicated genome regions, long-read sequencing technologies such as Pacific Biosciences (PacBio) and Oxford Nanopore provide an opportunity to overcome those limitations. In addition, Hi-C contact maps and Bionano optical mapping are scaffolding technologies that can bridge gaps between the sequence scaffolds and contigs in the final assembly to form chromosome level super-scaffolds as has been recently shown very successfully in the assembly of the goat genome (Bickhart et al. 2017).

Advances in long-reads DNA sequencing technologies, together with the development of new assembly algorithms and pipelines, have also enabled animal genomics researchers to go beyond a single reference assembly per species to compare across breeds and varieties within a single species. Such a “pan-genome” approach, by sampling a diverse set of individuals, would include all genomic segments that can be identified in representatives within the species and will provide a more complete resource for identifying and characterizing different types of genotype and genomic variations among individuals or distinct groups within a species (Sherman and Salzberg 2020; Gao et al. 2019). For rainbow trout, both the genome assemblies available in NCBI were developed from the DNA sequences of the Swanson rainbow trout doubled haploid (DH) homozygous line, one of 12 genetically diversified DH lines developed and maintained in Washington State University (Gao et al. 2018; Palti et al. 2014). The Swanson DH line is a YY male line developed from a semi-domesticated strain originated from the Kenai Peninsula, Alaska. The Arlee line is also a YY male line but in contrast it was originated from a domesticated hatchery strain that was originally collected from the northern California coast and the number of chromosomes between the two lines is also different. Although the number of chromosome arms in rainbow trout is constant, the number of chromosomes in a karyotype can vary from 2N=58 to 2N=64 due to centric fusions and fissions (Thorgaard 1983; Pearse et al. 2019). The karyotype of the Swanson line has 2N=58 with 29 haploid chromosomes (Phillips et al. 2006) and the Arlee karyotype includes 32 haploid chromosomes (2N=64) (Ristow et al. 1998). More evidence for the expected genetic and genomic differences between the two lines is shown in a phylogenetic analysis that was conducted using data from over 5,000 RAD SNPs where the Swanson and Arlee lines represented two distinct branches among the 12 lines (Palti et al. 2014).

The rainbow trout genome contains large polymorphic inversions on chromosomes Omy05 and Omy20 (Pearse et al. 2019). Recently, a second *de-novo* assembly for Atlantic cod has been useful for characterizing known inversions and identifying additional smaller inversions in the cod genome (Kirubakaran et al. 2020). By generating a second de-novo assembly for rainbow trout from a distinctly different genetic line we aim to further characterize the two known inversions in the rainbow trout genome and identify additional putative large-scale structural variants between the Swanson and Arlee lines. More broadly, multiple de-novo assemblies can be very useful in identifying and characterizing smaller structural genome variants as has been recently shown in plants (Editorial 2020; Gao et al. 2019).

In this report we generated a *de-novo* genome assembly for the Arlee homozygous line of rainbow trout to enable identification and characterization of genomic variants towards developing a rainbow trout pan-genome reference. The new assembly was generated using the PacBio long-read sequencing technology and scaffolding with Hi-C contact maps and Bionano optical mapping. A much-improved genome assembly was obtained, with the contig and scaffold N50 over 15.6 Mb and 39 Mb, respectively, and 95% of the assembly is anchored in chromosome sequences. The assembly is composed primarily of 32 major scaffolds corresponding to the karyotype of the Arlee line (2N=64). Through annotation and characterization of the duplicated genome loci that harbor the IGH gene family on chromosomes Omy12 and Omy13 (Magadan et al. 2019) we demonstrated the improved utility of this new assembly for annotation of some of the more complex areas of the genome.

## Materials and Methods

### DNA extraction and sequencing

DNA from a single fish from the Arlee DH line was extracted using the Qiagen DNeasy kit from a fresh blood sample collected in EDTA and kept on ice until DNA extraction within 24 hours to ensure yield of high molecular weight DNA for long-reads sequencing. The DNA samples was sequenced at the core facility of the University of Delaware (Newark, Delaware, USA) using the PacBio RS-II system which generated approximately 268 Gb of sequence data (111x coverage). The PacBio reads length N50 was 33 kb and the average read length was 20 kb. For sequencing errors corrections and polishing the genome assembly, 1.6 billion paired-end reads (2 x150bp) of Illumina short-reads were generated by Admera Health (South Plainfield, NJ, USA) from the same Arlee DNA sample. The short-read sequences were processed with the program cutadapt to remove the Illumina sequencing adaptors leaving 1.59 billion reads with total length of 477 Gb (∼199x genome coverage).

### Construction of the assembly contigs

The initial assembly of sequence contigs was generated from the PacBio long-reads using the Canu (Version 1.8) assembler (Koren et al. 2017) with the options of correctedErrorRate=0.085, corMhapSensitivity=normal, and ovlMerDistinct=0.975. After the “correction” and “trimming” steps, Canu retained 2.1 million high quality reads with 87.5 Gb (∼36x) total length from the raw PacBio reads. The retained reads were then assembled into 1,591 contigs in the final step of Canu. Next, two rounds of “arrow” polishing were conducted on the contigs. In each round, the PacBio raw reads were first mapped to the contigs using the minimap2 (Version 2.15-r915) aligner (Li 2018), and then, based on the alignment results, the contigs were polished using the PacBio “arrow” algorithm (https://github.com/PacificBiosciences/GenomicConsensus). To further fix the indel errors in the contigs, the Illumina short-read data were aligned to the contigs using the BWA-MEM (Version 0.7.17-r188) aligner (Li 2013) with the default parameters and indel variants were called using the freebayes (Version 1.0.0) program (Garrison and Marth 2012) based on the alignments without mismatch (using the option -- read-mismatch-limit 0). Finally, the contigs were corrected based on the called indel variants using the bcftools program (Li et al. 2009).

### Construction of the assembly scaffolds and chromosome sequences

To improve the contiguity of the assembly, we first used Bionano optical mapping with the Saphyr platform (Bionano Genomics, San Diego, CA, USA) to scaffold the assembly contigs. High molecular weight DNA were extracted in agarose gel plugs from fresh blood of the same Arlee DH line fish by Amplicon Express (Pullman, Washington) and the extracted DNA molecules were labeled with Direct Label Enzyme (DLE-1). A total of 11 million DNA molecules were generated with 1.2 Tb total length (average length 111 Kb). These raw mapping data were processed with the Bionano Solve software (Solve3.3_10252018) (https://bionanogenomics.com/support-page/bionano-access-software/) to generate a de novo Bionano assembly guided by the contigs of the polished PacBio assembly. Two million molecules with N50 of ∼224 Kb (∼200x genome coverage) and average label density of 11.02 sites per 100 Kb were selected from the raw data after the filtering step, and 905 Bionano genome maps were generated with a total length of ∼2.29 Gb and N50 of ∼16.3 Mb. The Bionano genome maps and the polished Canu assembled contigs were then scaffolded using the Bionano hybrid scaffold pipeline. This resulted in the joining of 664 contigs (∼2.26 Gb) into 345 scaffolds and breaking of 13 Canu contigs that were considered by the Bionanao pipeline to be chimeric joins. In the next step the scaffolds were manually checked with the alignments of the Bionano genome maps and the Canu contigs to the hybrid scaffolds, and one scaffold was found to be incorrect and was disassembled. In addition, sequence duplications that were artificially generated by the Bionano assembly software were removed from 88 scaffolds. These duplicated sequences were found on both sides of the 13 bp gaps that were inserted by the Bionano software to indicate the presence of small gaps of undetermined size or undetermined flanking sequence overlaps between pairs of Canu contigs that were joined by the Bionano hybrid scaffolds pipeline.

The Hi-C proximity ligation sequence data was used to generate chromosome scale scaffolds from the Bionano scaffolds assembly. Fresh blood (100 ul) was collected in ethanol (750 ul) and immediately placed and then shipped on dry ice. The Hi-C library was generated and sequenced to produce 857 million Illumina paired-end reads (2×150 bp) (Arima Genomics, San Diego, CA, USA). The reads were aligned to the Bionano scaffold and the alignments were filtered following the Arima-HiC mapping pipeline (http://github.com/ArimaGenomics/mapping_pipeline). Briefly, sequence reads from each end were mapped separately to the Bionano hybrid scaffolds using the BWA-MEM aligner (Li 2013). The chimeric 3’ side of the aligned reads that crossed the ligation junctions was removed and the reads that were mapped from both ends with the mapping quality score higher than 10 were retained. PCR duplicates in the Hi-C sequences were identified and removed using the Picard Toolkit (http://broadinstitute.github.io/picard/). After these filtering steps, 203 million paired reads alignments were retained, including 61 million inter-scaffold alignments. The paired-read alignment data were then processed with the program SALSA (Ghurye et al. 2019) which broke 65 chimeric scaffold and contig sequences and merged 287 sequences into 97 Hi-C scaffolds.

Linkage information from two single nucleotide polymorphism (SNP) genetic maps was used to generate chromosome sequences from the Hi-C assembly scaffolds. The program NovoAlign (http://www.novocraft.com/products/novoalign/) was used to map the flanking sequences of each SNP marker to the scaffolds and the genetic linkage information was used to anchor the scaffolds to chromosomes and then to order, orient and concatenate the scaffolds into 29 chromosomes. With the guidance of the linkage information, 13 Hi-C scaffolds were broken into 32 smaller scaffolds at 19 Hi-C join sites due to incorrect ordering or wrong orientation of the joined scaffolds, or due to the insertion of a smaller scaffold at the join site of the larger scaffold. The two linkage maps were previously used for the same purpose in the OmyK_1.0 chromosome sequences assembly (Pearse et al. 2019). The primary map was generated with genotypes from the 57K high-density SNP chip (Palti et al. 2015). With 29 linkage groups, this genetic map contained 46,174 SNP markers and was based on genotype data from a pedigree of 5,716 individual fish from 146 full-sib families. The second genetic map was generated from 9,551 restriction site associated DNA (RAD) SNP markers with genotype data from 10 full-sib families from the USDA-NCCCWA broodstock (Vallejo et al. 2017), and was used to anchor additional scaffolds that could not be anchored with the primary map to chromosomes.

### Detection and quantification of repetitive elements

RepeatModeler (Version 2.0.1) (http://www.repeatmasker.org) with RECON (Version 1.08) (Bao and Eddy 2002) and RepeatScout (Version 1.0.6) (Price et al. 2005) were used for de novo identification and modeling of transposable elements (TEs) in this genome assembly. This generated a repeat library with 2,492 sequences, each representing a consensus sequence of a TE family. Using this new repeat library, 48.28% of the assembly was masked with RepeatMasker (Version 4.0.6). Detailed information on the repeat masking results and the repeat library sequences can be found in **Supplementary File 1**.

To compare and align chromosome sequences between genome assemblies repeat masked sequences were aligned using megablast with the parameters -evalue 1.0e-100 and -word_size 50.

### Mapping and annotation of the IGH duplicated locus

Chromosomes OmykA12 and OmykA13 from the Arlee clonal line genome assembly (USDA_OmykA_1.1) were analyzed to locate immunoglobulin heavy chain (IGH) loci. Previously published IGH constant (IGHC) exon sequences from rainbow trout and Atlantic salmon (Magadan et al. 2019; Yasuike et al. 2010; Solem et al. 2001) were used as queries to identify the chromosomal regions containing immunoglobulin genes. The IGHC exon sequences search was performed with the available BLAST tool at the Galaxy website (www.usegalaxy.org) and the detailed analysis and visualization with the program SnapGene Viewer (www.snapgene.com).

Comparative phylogenetic studies were carried out with MEGA7 (Kumar et al. 2008; Tamura et al. 2013), by using the ClustalW and Muscle algorithms to perform alignments. Subsequently, the Neighbour-Joining and Maximum Likelihood methods were used to plot phylogenetic trees (pair-wise deletion, JTT or WGA matrix). The veracity of these trees was studied using the above-mentioned method and by executing 3,000 replicates of bootstrapping.

## Data availability

All raw sequence data is available in NCBI BioProject number PRJNA639144. The genome assembly is available as GenBank accession GCA_013265735.3. The RefSeq assembly annotation is available as accession GCF_013265735.2.

## Results and Discussion

### Genome assembly

The rainbow trout genome assembly presented in this paper was based on DNA from the Arlee homozygous line and was developed using PacBio long-reads sequencing technology. The PacBio sequences were assembled into contigs with the Canu program and those contigs were joined into scaffolds using Bionano optical mapping and Hi-C proximity ligation sequence data, also known as proximity or contact mapping. Using genetic linkage information, the scaffolds from the Hi-C assembly were then anchored to and ordered on chromosomes to generate the chromosome sequences of the USDA_OmykA_1.1 assembly (GCA_013265735.3).

The Canu sequence assembly was composed of 1,591 contigs with a total length of ∼2.34 Gb, N50 of 9.8 Mb, and BUSCO score of 95.9% (using actinopterygii_odb9 as the lineage dataset). After the polishing steps with Arrow and Freebayes the BUSCO score was improved to 96.2%. More detailed statistics for each step in the assembly pipeline are shown in Table 1. The Bionano hybrid scaffolding pipeline used the optical map to join neighboring or overlapping Canu contigs and to break chimeric sequence contigs that were misassembled by Canu. The final Bionano scaffolded assembly included 344 scaffolds of joined Canu contigs and 700 contigs that could not be joined into scaffolds after removing 254 contigs from the original Canu assembly that were shorter than 15 Kb. The N50 of these 1,044 scaffolds and contigs in the Bionano assembly is 28 Mb (**Table 1**). The Hi-C ligation proximity mapping broke 65 chimeric scaffold and contig sequences from the Bionano assembly and merged 287 sequences into 97 Hi-C scaffolds. Including the 822 sequences that were not joined by Hi-C scaffolding, the Hi-C scaffolded assembly is composed of 919 scaffolds with N50 of 47.5 Mb (Table 1). A list of the 919 Hi-C assembly scaffolds sorted by sequence length is presented in **Supplementary File 2**. Using linkage information from previously published genetic maps (Pearse et al. 2019), 209 Hi-C scaffolds were anchored to 29 linkage groups to generate 29 chromosome sequences in a total length of ∼2.23 Gb. The total length of the 710 unplaced scaffolds that are not anchored to a chromosome is ∼108 Mb.

**Table 1.**
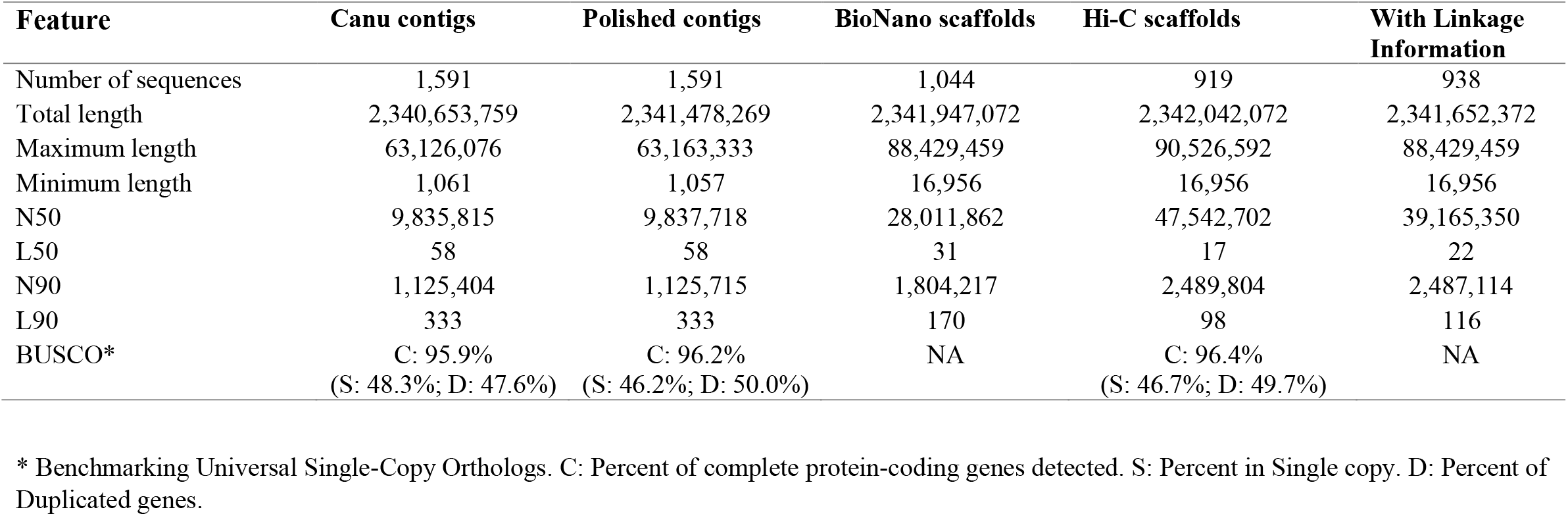
Statistical data summary of the USDA_OmykA_1.1 (Arlee) rainbow trout genome assembly.

| **Feature** | **Canu contigs** | **Polished contigs** | **BioNano scaffolds** | **Hi-C scaffolds** | **With Linkage Information** |
| --- | --- | --- | --- | --- | --- |
| Number of sequences | 1,591 | 1,591 | 1,044 | 919 | 938 |
| Total length | 2,340,653,759 | 2,341,478,269 | 2,341,947,072 | 2,342,042,072 | 2,341,652,372 |
| Maximum length | 63,126,076 | 63,163,333 | 88,429,459 | 90,526,592 | 88,429,459 |
| Minimum length | 1,061 | 1,057 | 16,956 | 16,956 | 16,956 |
| N50 | 9,835,815 | 9,837,718 | 28,011,862 | 47,542,702 | 39,165,350 |
| L50 | 58 | 58 | 31 | 17 | 22 |
| N90 | 1,125,404 | 1,125,715 | 1,804,217 | 2,489,804 | 2,487,114 |
| L90 | 333 | 333 | 170 | 98 | 116 |
| BUSCO* | C: 95.9% | C: 96.2% | NA | C: 96.4% | NA |
|  | (S: 48.3%; D: 47.6%) | (S: 46.2%; D: 50.0%) |  | (S: 46.7%; D: 49.7%) |  |

\* Benchmarking Universal Single-Copy Orthologs. C: Percent of complete protein-coding genes detected. S: Percent in Single copy. D: Percent of Duplicated genes.

### 32 Major Hi-C scaffolds match the Arlee karyotype

The longest 32 Hi-C scaffolds are highlighted in **Supplementary File 2**. The graphic presentation of the Hi-C contact map generated by Juicebox (Durand et al. 2016) is shown in **Supplementary File 3** and also highlights the presence of 32 major scaffolds in the genome assembly. Those 32 scaffolds are substantially longer than the rest of the genome assembly scaffolds. Their lengths are in the range of 33.3 Mb to 90.5 Mb, which is close to the length range of the chromosome sequences, and the total length of these 32 scaffolds is ∼81% of the total length of the chromosome sequences assembly. Of those 32 major assembly scaffolds, 26 were mapped in a one to one ratio to 26 linkage groups aligning directly to 26 chromosomes from the Swanson assembly. The other six were mapped in pairs to the remaining three linkage groups, and one of each pair can be aligned perfectly to one of the two chromosome arms from the metacentric Swanson assembly chromosomes that are associated with the same three linkage groups. The lengths of each of the 32 major scaffolds and the chromosome or chromosome arm to which they were mapped are displayed in **Figure 1** and listed in **Supplementary File 4**. The three chromosomes from the Swanson assembly that can be aligned with two major scaffolds each are Omy04, Omy14, and Omy25. Those three chromosomes have been previously shown to be associated with variable chromosome numbers due to centric fusions or fissions in rainbow trout (Pearse et al. 2019). Plots of the genetic distance from the linkage genetic maps versus the physical distance from this Arlee genome assembly for each of the three chromosomes are shown in **Figures 2a-2c**. The gap in the linkage maps at the centromeric region in each chromosome plot is caused by a fission splitting the metacentric chromosome from the Swanson genome assembly into two acrocentric chromosomes in the Arlee genome assembly. As shown in Figure 2, those linkage map gaps correspond perfectly to the boundaries between a pair of two of the 32 major scaffolds. Each pair of scaffolds was uniquely mapped to one of those three chromosomes with each scaffold mapping uniquely to one of the two chromosome arms, indicating that the Arlee line haploid karyotype is composed 32 chromosomes. Our results from the alignment of the genome assembly to the linkage groups are in complete agreement with the known karyotype (2N=64) of the Arlee rainbow trout line (Ristow et al. 1998). To resolve the karyotype difference with the Swanson genome and remain consistent with the nomenclature of the Omyk_1.0 genome assembly, we named the Arlee chromosomes aligned with the Swanson q chromosome arms with the same chromosome number and assigned new numbers to the p arms according to their physical length. Hence the scaffolds mapped to the linkage groups corresponding to the Swanson Omy04p, Omy25p and Omy14p were renamed chromosomes OmyA30, OmyA31 and OmyA32, respectively.

**Figure 1:**
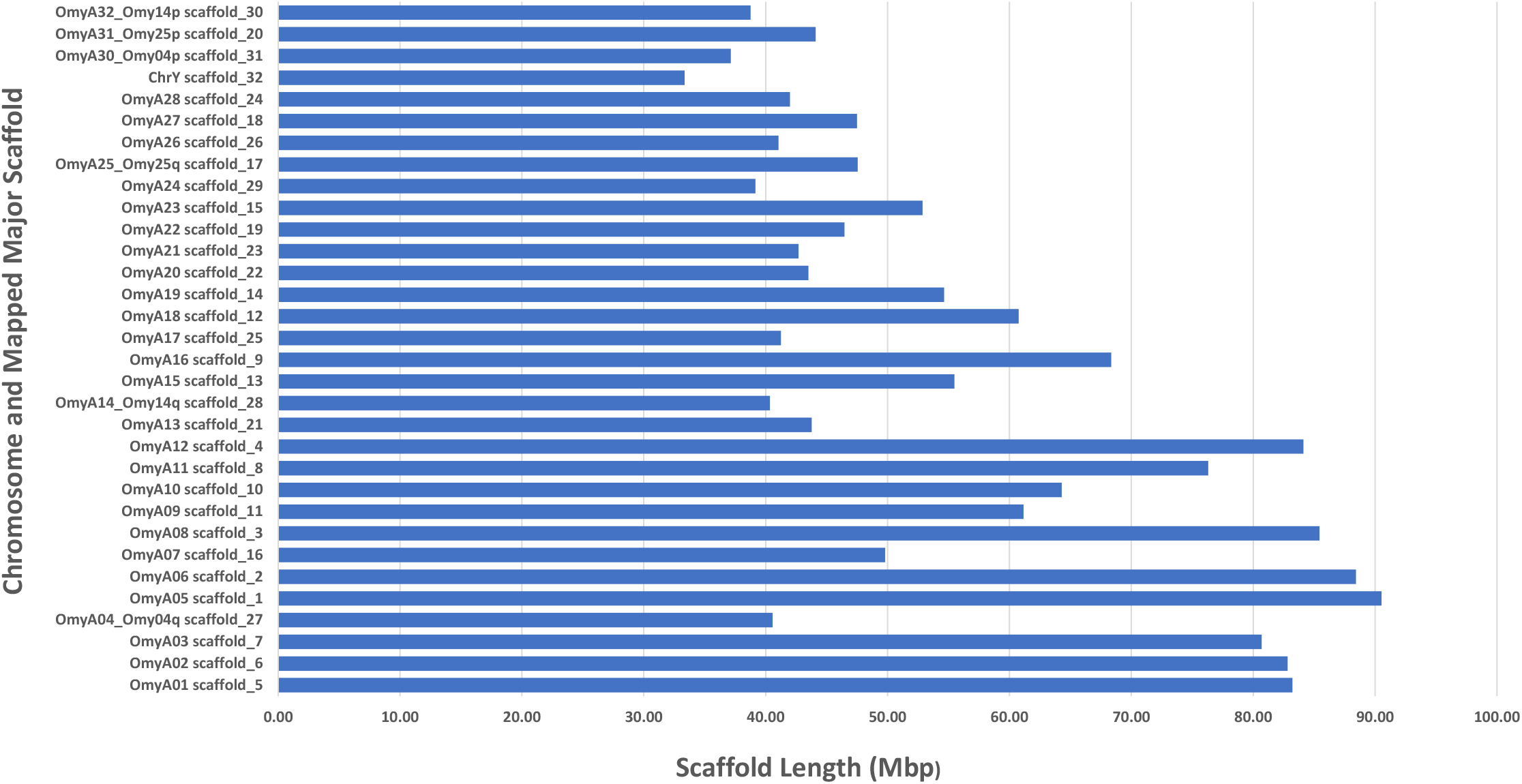
Length distribution of the 32 major Hi-C scaffolds and the chromosome number to which each scaffold was mapped in the rainbow trout Arlee line genome assembly.

**Figure 2:**
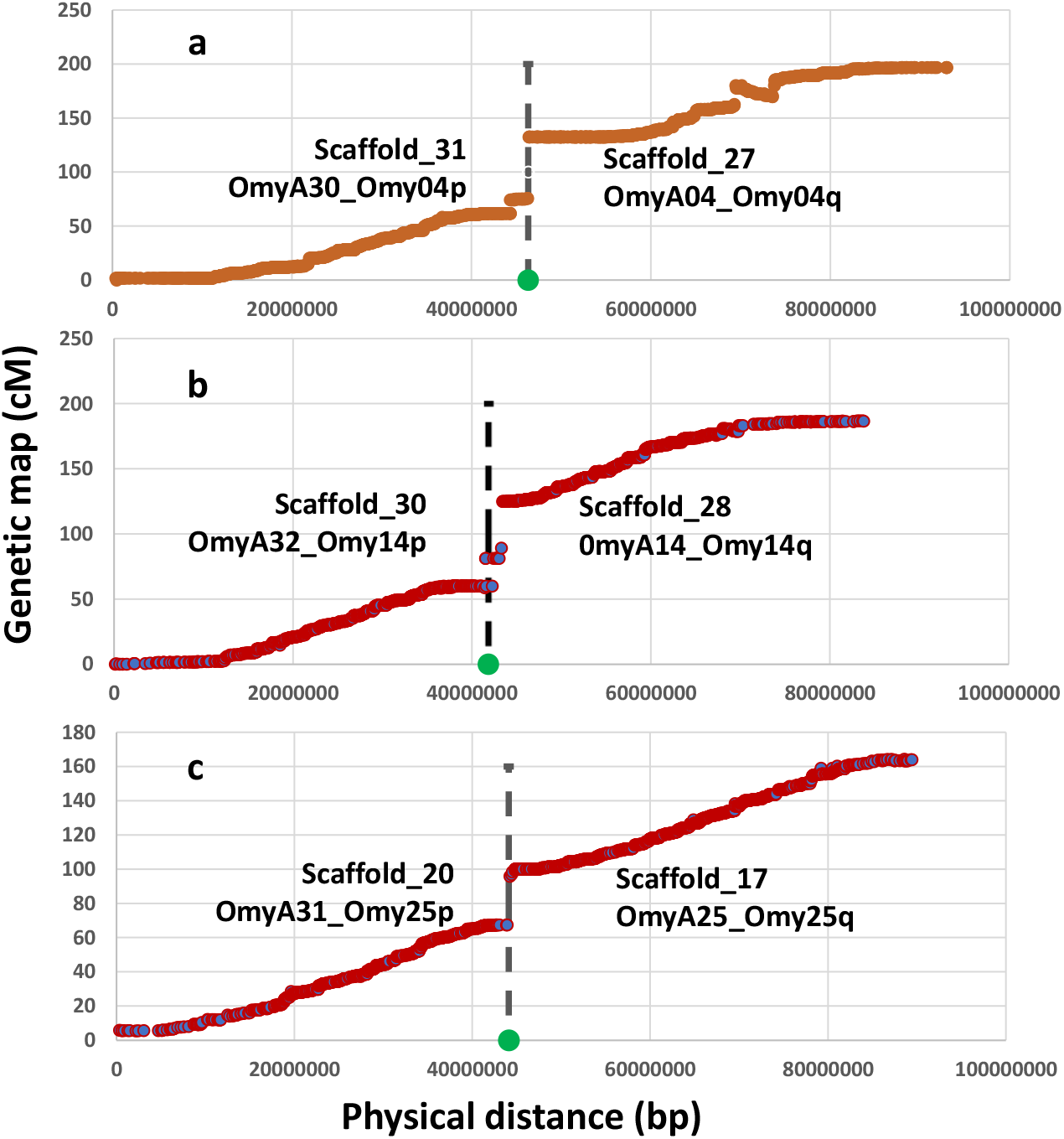
Genetic distance from the female genetic linkage map (Pearse et al., 2019) plotted on the Y axis versus physical distance on the X axis for the Arlee assembly chromosomes OmyA04/OmyA30 (a), OmyA14/OmyA32 (b), and OmyA25/OmyA31 (c) with the corresponding chromosome arms from the Swanson assembly showing the location of centric fusions or fissions in those three chromosomes. The vertical dash line and green dot in each plot mark the boundary between the two major scaffolds forming the corresponding chromosome sequences in the Arlee genome assembly.

Approximately 2.23 Gb of scaffolded sequence, or 95% of the total Arlee assembly length, was anchored through alignment of SNPs flanking sequences to linkage groups and were included in the 32 chromosome sequences. The length of each chromosome sequence and the number of Hi-C scaffolds per chromosome are shown in **Figure 3** and are listed in **Supplementary File 4**. The lengths of the chromosome sequences ranged from 41.8 Mb to 103.8 Mb and the number of Hi-C scaffolds per chromosome ranged from 1 to 17, with an average of 7 scaffolds. Considering that 13 of the original Hi-C scaffolds were broken into 32 smaller scaffolds to address ordering and orientation errors based on the linkage map information, the final genome assembly is composed of 228 scaffolds that were anchored to 32 chromosomes and 710 unplaced scaffolds.

**Figure 3:**
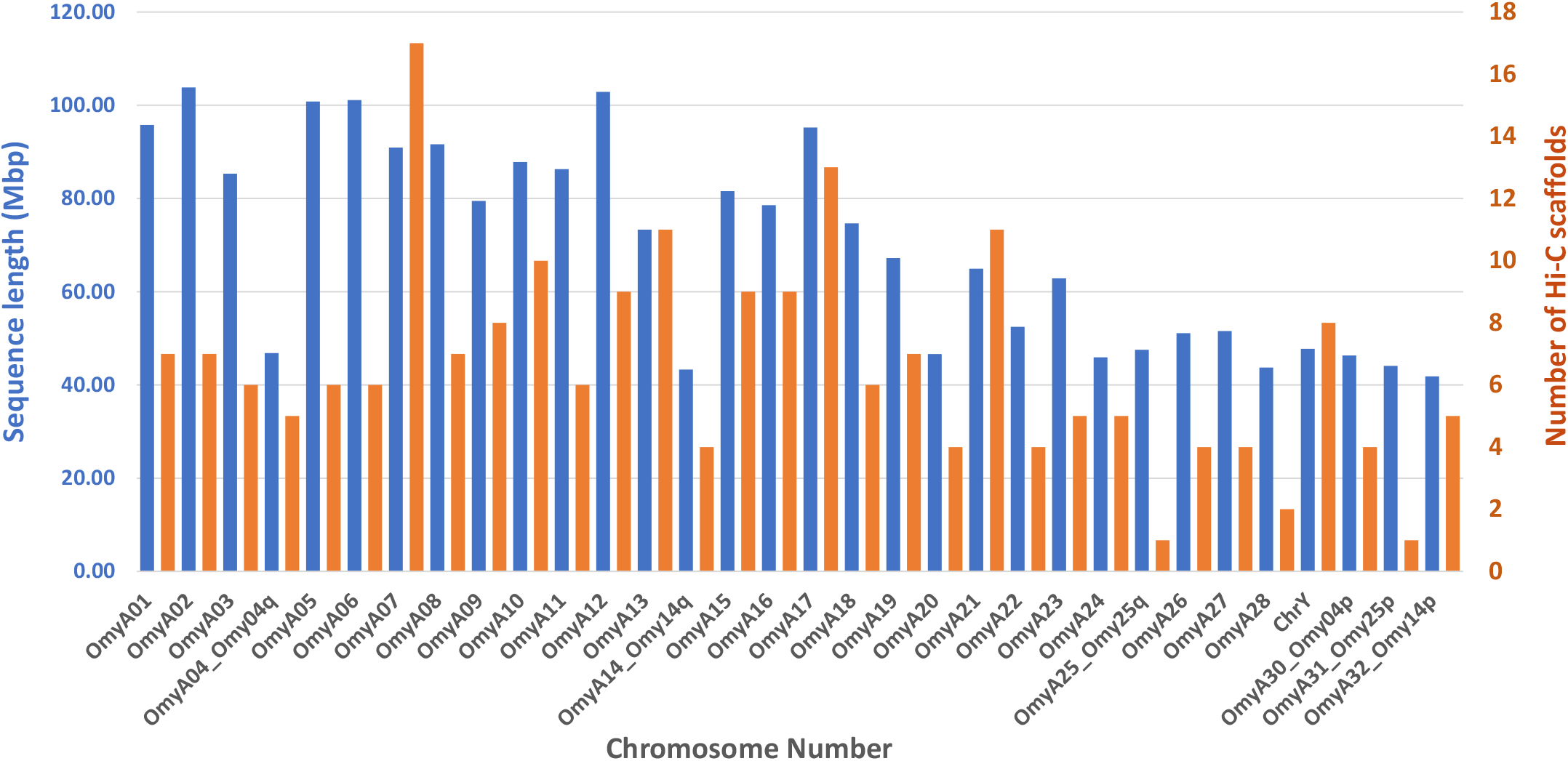
Length (in blue) and number of scaffolds (in orange) for each chromosome in the rainbow trout Arlee line genome assembly.

This Arlee genome assembly has greater contiguity than the other two publicly available rainbow trout genome short-reads assemblies (Berthelot et al. 2014; Pearse et al. 2019). It contains only 486 gaps compared to 420,055 spanned and 7,839 unspanned gaps in the Omyk_1.0 assembly which was derived from the Swanson line (Pearse et al. 2019). There are 438 gaps in the Arlee chromosome sequences with an average of 14 per chromosome (**Supplementary File 5**). The total length of the new assembly is 2.34 Gb, which is very close to the 2.4 Gb estimated size of the rainbow trout genome (Hardie and Hebert 2003). The overall score of complete protein-coding genes detected with the Benchmarking Universal Single-Copy Orthologs (BUSCO) assessment was 96.4% compared to 91.1% detected in the Omyk_1.0 assembly, which means an increase by 5.5% in the detection of protein-coding genes that are expected to be found in the trout genome. Furthermore, 51% of the complete BUSCO protein coding ortholog genes detected in the current Arlee assembly are duplicated in the genome, compared to 43% that are duplicated in the Omyk_1.0 assembly. The expectation of ∼50% duplicated genes in the genome is consistent with previous results from screening the whole-genome BAC library with probes from protein-coding genes (Palti et al. 2004), and hence provides another indication that the current Arlee assembly provides a more complete reference genome for rainbow trout.

### Chromosomal inversions

The rainbow trout genome has been previously shown to contain large polymorphic inversions on chromosomes Omy05 and Omy20 (Pearse et al. 2019). The double inversion on Omy05 is composed of two adjacent inversions of approximately 55 Mb that are thought to act as a supergene complex associated with adaptive traits such as migratory behavior (Pearse et al. 2019) and hatching time (Miller et al. 2012). The two alternative inversion karyotypes on Omy05 were categorized as ancestral (A) and rearranged (R) based on the sequence and structural synteny to the genomes of other salmonids such as Atlantic salmon, Coho salmon, and Arctic char (Pearse et al. 2019). Using a haplotype composed of 475 individual SNP genotypes from the Axiom 57K SNP chip (Palti et al. 2015) it was shown that the Swanson Omy05 inversion karyotype is RR and the Arlee’s is AA (**Supplementary File 5**). However, alignment of the Omy05 chromosome sequences between the OmykA_1.1 (Arlee) and Omyk_1.0 (Swanson) assemblies as shown in **Figure 4a** revealed that there are no inversions between them. The cause for this is an error in the assembly of the Omy05 chromosome sequence in the Omyk_1.0 assembly that was guided by a linkage map predominantly from fish with the A karyotype (Pearse et al. 2019). This error must be corrected in a revision of the Swanson line genome assembly which we plan to generate using long-reads sequencing technology.

**Figure 4:**
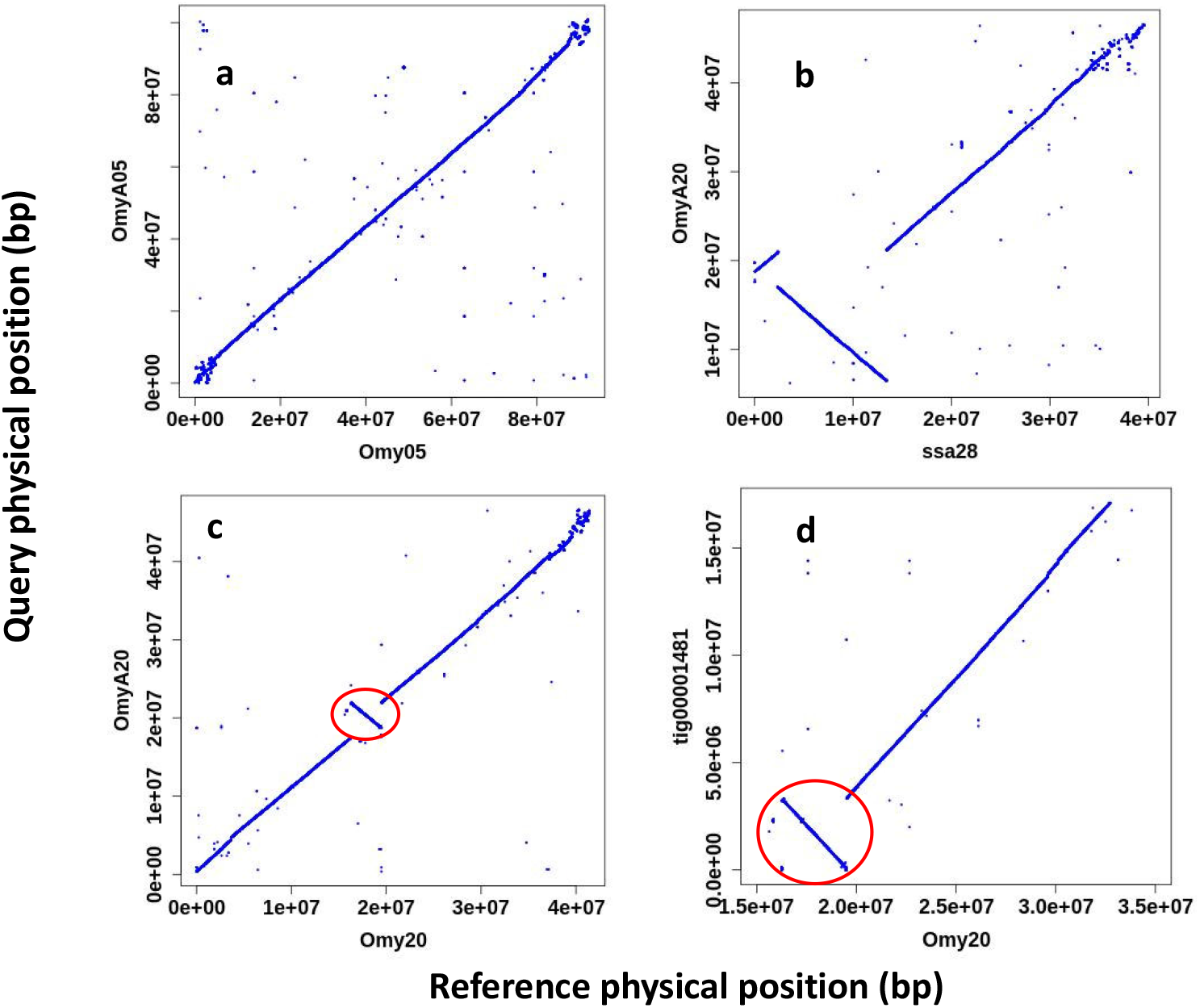
Alignment plots of large inversion regions on chromosomes 5 and 20 in the rainbow trout genome. a. OmyA05 from this Arlee genome assembly aligned to Omy05 and showing that as oriented in the current Swanson assembly (Pearse et al., 2019) both are arranged according to the Ancestral (A) karyotype of the double inversion on chromosome 5. b. OmyA20 aligned to chromosome Ssa28 of the Atlantic salmon genome assembly (Lien et al., 2016) showing the major inversion and indicating that the Arlee genome has the Rearranged (R) karyotype for the inversion on rainbow trout chromosome 20. c. OmyA20 aligned to Omy20 of the current Swanson assembly and showing only a small inversion close to the center of the chromosome while the major inversion is aligned in the same orientation between the two rainbow trout genome assemblies. d. Canu contig tig00001481 of OmykA_1.1 assembly aligned to Omy20 of the current Swanson assembly and showing that the smaller inversion is within the boundary of the Arlle assembly contig, and hence not likely to be caused by scaffolding error in the Arlee assembly.

The inversion on Omy20 has not been shown yet to our knowledge to be associated with a known phenotypic diversity. The alternative genotypes could not be identified using SNP markers as was done with the Omy05 double-inversion karyotype, so the new Arlee de-novo assembly provided an opportunity to better define the Omy20 inversion karyotype. To characterize the Omy20 inversion karyotype in the Arlee genome, the repeat masked OmyA20 sequence was aligned to the repeat masked Atlantic salmon genome assembly (GCF_000233375.1) (Lien et al. 2016). The alignment of OmyA20 to the Atlantic salmon Ssa28 chromosome sequence (**Figure 4b**) indicates that there is an 11-Mb inversion between the two genomes, which is close to what was reported for the Omyk_1.0 assembly in this region. Hence, we demonstrate that the Arlee genome has the R inversion karyotype for chromosome 20. The alignment of OmyA20 to Omy20 of Omyk_1.0 (**Figure 4c**) is showing a smaller 3-Mb inversion between the two rainbow trout genome assemblies. However, such a small inversion between assemblies can be the result of a single smaller scaffold that was not oriented correctly in the chromosome sequence rather than a real structural genome variant between the two rainbow trout lines. Further analysis of the sequence in this region shows that this smaller 3-Mb inversion is within the boundaries of the 17 Mb Canu contig (tig00001481) in the major scaffold of OmyA20 (**Figure 4d**) and therefore was not caused by miss-orientation of a smaller contig or scaffold in the Arlee genome assembly.

Furthermore, throughout the genome we identified 15 such small inversions in the size of 1 to 5 Mb between the OmykA_1.0 (Arlee) and Omyk_1.0 (Swanson) assemblies that reside within the boundaries of Canu contigs from the Arlee assembly (**Supplementary File 6**). Upon improvement of the Swanson line genome assembly with long-reads sequencing technology we plan to repeat these comparative chromosome sequence alignments to assess if those smaller inversions appear to be the result of real genome variants rather than miss-orientation of small scaffolds from the short-reads genome assembly.

### Chromosome location of the sdY gene

The improved contiguity and near completeness of this assembly has substantially improved the mapping accuracy for the rainbow trout genome. For example, the sexually dimorphic on the Y chromosome protein (sdY) gene is the rainbow trout sex-determining gene (Yano et al. 2012), which is very important in analyzing sex-related traits. Therefore, mapping it accurately to a chromosome location and identifying the flanking DNA sequences is of great interest. However, in Omyk_1.0, this gene is in a 65.5 Kb unplaced scaffold (NW_018580030), and in AUL_PRJEB4421_v1 it is also in a 1.9 Kb unplaced scaffold (scaffold_11164). In both assemblies, there was no genetic marker present in either of those scaffolds to anchor and place the scaffold within a chromosome sequence. In Omyk_1.0 its scaffold location in chromosome Omy29 was estimated using indirect indicators, including the between-sexes F_ST_ data in a group of rainbow trout and the alignment of the bacterial artificial chromosome (BAC) sequence containing the sdy gene to the genome assembly (Pearse et al. 2019). Here in the Arlee line genome assembly, the sdy gene is found in scaffold_81, which is a 3-Mb sequence containing 24 SNP markers from the genetic map that were used to anchor the scaffold to chromosome Y (CM023247.2; OmyA29) and place it between nucleotide positions 2,936,690 bp and 6,038,050 bp.

### Mapping and annotation of the duplicated IGH Loci

The new assembly of the rainbow trout genome aided by data from long-reads sequencing technology offers the opportunity to improve the gene annotation of complex loci such as the immunoglobulin heavy chain (IGH) locus. As was previously identified in the reference genome from the Swanson line (Magadan et al. 2019), the rainbow trout IGH genes are located within two regions, IGHA and IGHB, on chromosomes 13 and 12, respectively. Both loci, IGHA and IGHB, follow a pattern of translocon configuration, with a number of IGHV genes followed by units comprising several IGHD and IGHJ gene segments, and a region comprising IGHC genes. In salmonids, IGHC genes belong to three subgroups IGHM, IGHD and IGHT. When functional, the C-REGION of the heavy chain defines these three isotypes; IG-Heavy-Mu (heavy chain of the IgM class), IG-Heavy-Delta (heavy chain of the IgD class) and IG-Heavy-Tau (heavy chain of the IgT class).

In the Arlee genome assembly, the IGHA locus spans ∼500 kb (from 51,600,000 to 52,000,000 bp), and the IGHB locus ∼450 kb (from 84,365,000 to 84,810,000 bp).The number of identified IGHC genes is higher than those previously annotated in the Swanson reference genome (**Figure 5**), and the organization of their corresponding CH exons is more consistent with the mRNA expression data in the literature (Hansen et al. 2005; Zhang et al. 2017). A single Cμ gene, three Cδ and three Cτ genes were identified in the Arlee-line assembly IGHA and IGHB loci **(Figure 5**). In both Arlee and Swanson, the two duplicated IGH loci present a rather similar gene cluster containing Cμ-Cδ genes (i.e. IGHM-IGHD) at the 3’end of the IGH loci, all of them in forward orientation. However, there are differences in the length and CH exon composition of the IGHD genes identified in the Arlee IGHA locus (Chr.13). In both assemblies there are 4 IGHC exons coding for IgM, while the number of IGHC exons coding for IgD is 10 in the Arlee genome and 11 in the Swanson genome assembly (**Figure 5**).

**Figure 5:**
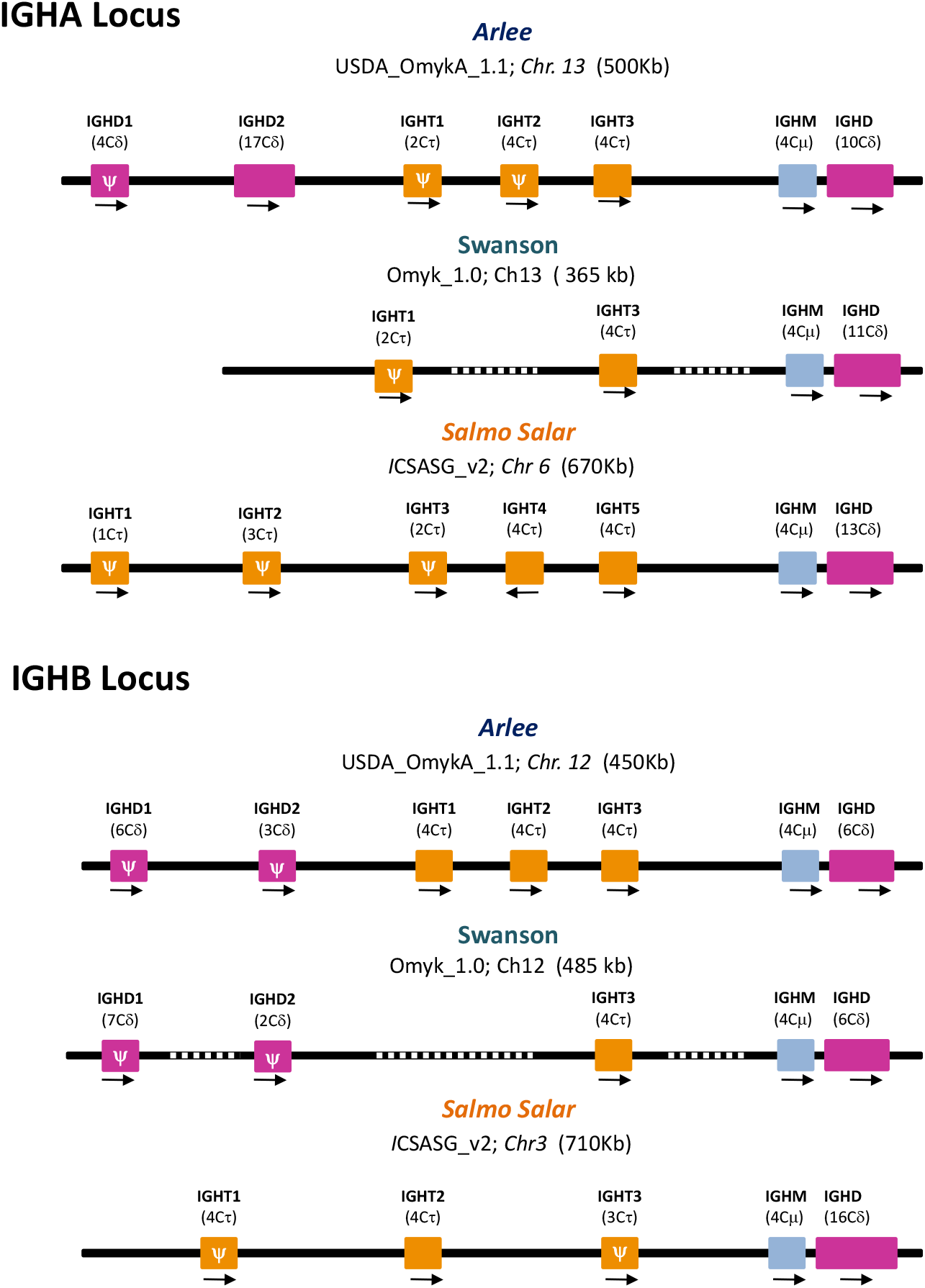
The overall configuration of IGHC genes identified in the rainbow trout genome assemblies USDA_OmykA_1.1 (Arlee strain) and Omyk_1.0 (Swanson strain). Rainbow trout IGH loci on chromosome 13 (IGHA) and on chromosome 12 (IGHB) are shown in comparison to IGHC genes that were previously characterized in *Salmo salar* (ICSASG_v2). The boxes represent the IGHM, IGHD and IGHT genes annotated. The number of CH exons are indicated between brackets and the transcriptional direction is shown by arrowheads. Pseudogenes are represented by (Ψ). Dashed line indicates sequence gap. IGH maps are not to scale.

The genome organization of the teleost IGHD genes differs between fish species (Saha et al. 2004; Gambón-Deza et al. 2010; Hirono et al. 2003; Hansen et al. 2005). In Atlantic salmon, the IGHD gene in the IGHA locus, which is situated downstream of the IGHM gene, is arranged in this configuration: Cδ1-(Cδ2-Cδ3-Cδ4)_3_ –Cδ5-Cδ6-Cδ7-TM1TM2 (**Figure S1**). In contrast, the phylogenetic analysis of Cδ exons found in the Arlee and Swanson genome assemblies suggests a different composition and organization in rainbow trout: Cδ1-(Cδ2-Cδ3-Cδ4)2 – Cδ2 - Cδ7 – TM1TM2 (**Figures S1 and S2**), lacking Cδ5 and Cδ6 exons and a duplication of Cδ2-Cδ3-Cδ4. Even more, the new Arlee genome assembly allows correction of likely artefacts in the Omyk_1.0 genome assembly that have generated aberrant localizations and duplications of Cδ exons (Cδ2, Cδ3 and Cδ4; **Figure S1**). The alignment of known sequences of the rainbow trout secreted IgD with the deduced amino acid sequences of 10 Cδ exons from the Arlee genome assembly also supports this Cδ exon configuration (**Figure S3**).

Two additional IGHD genes, located upstream of the IGHM-IGHD cluster, were identified in the Arlee assembly IGHA locus (**Figure S1**). IGHD1 contains four Cδ exons, and IGHD2 presents 17 Cδ exons (one of them a pseudogene). The sequence comparison with *S. salar* Cδ exons, and the phylogenetic analysis indicates that the IGHD2 gene follows this configuration: Cδ1-(Cδ2-Cδ3-Cδ4)_5_ – Cδ2, and the IGHD1 gene is in this configuration: Cδ4-Cδ5-Cδ6-Cδ7. Interestingly, IGHD1 presents two exons that nicely cluster with S. salar Cδ5 and Cδ6 (**Figure S4**). The absence of IGHJ –IGHD segments or IGHM gene linked to IGHD1 or IGHD2, and the presence of stop codons in IGHD1, suggest that both genes are not functional, although this must be confirmed through RNA expression and other functional analyses. Additional IGHD genes at the 5’end of the IGHB locus can be also identified in the Arlee and Swanson assemblies, with similar number of CH exons and similar presence of stop codons (**Figure 5**).

Previous studies performed in rainbow trout have reported 3 IgT subclasses based on gene expression analysis (Zhang et al. 2017). However, in the Swanson Omyk_1.0 genome assembly only two functional IGHT genes were identified (Magadan et al. 2019), one in each IGH locus (**Figure 5**). In the Arlee genome assembly (USDA_OmykA_1.1), we identified three IGHT genes in the IGHA locus, of which only one is functional, and three functional IGHT genes in the IGHB locus. The annotation of only one of the three functional IGHT genes in the Swanson-IGHB locus is likely caused by the presence of a big gap in the same region that interrupts the open reading frames annotation. The alignment and similarity analysis of amino acid IgT sequences deduced from the four functional Arlee-IGHT genes indicate that IGHT1 and IGHT3 from the Arlee IGHB are identical (**Figure S5A**). The comparison with the constant region of previously identified rainbow trout IgT subclasses suggest that these IGHT1 and IGHT3 genes codify for IgT2 subclass, while IGHT1 from the IGHA locus codifies for the IgT1 subclass (**Figure S5B**). However, the identity between the IGHT3 from the Arlee IGHB locus and the IgT3 subclass identified by Zhang et al. (2017) is not high (< 85%). Further functional studies will be needed to evaluate the different roles of the functional IGHT genes identified in this new Arlee-line genome assembly.

### Genome annotation

The new Arlee genome assembly was annotated with the RefSeq annotation pipeline in NCBI (*Onchorhynchus mykiss* Annotation Release 101; GCF_013265735.2). A summary of the RefSeq annotation statistics in comparison with the Swanson assembly annotation in RefSeq (Release 100; GCF_002163495.1) is shown in Table 2. The number of predicted protein-coding genes is similar between the two assemblies with 41,896 vs. 42,884 in the Arlee and Swanson assemblies, respectively. However, the number of predicted non-coding genes is substantially higher with 28,007 vs. 10,499 in the Arlee and Swanson assemblies, respectively. Similarly, the numbers of coding sequences, exons and introns are substantially higher in the Arlee assembly due to its improved contiguity and near-completeness.

**Table 2.** Comparative statistical data summary of the Refseq genome annotation for the USDA_OmykA_1.1 (Arlee) and the Omyk_1.0 (Swanson) rainbow trout genome assemblies.

| **Feature** | **Arlee Assembly**<br>**(USDA_OmykA_1.1)** | **Swanson Assembly**<br>**(Omyk_1.0)** |
| --- | --- | --- |
| Genes | 69,903 | 53,383 |
| Protein-Coding Genes | 41,896 | 42,884 |
| Non-Coding Genes | 28,007 | 10,499 |
| Average Genes Length | 18,975 bp | 19,485 bp |
| Median Length | 4,422 bp | 7,247 bp |
| Min Length | 53 bp | 66 bp |
| Max Length | 1,190,608 bp | 1,178,930 bp |
| Pseudogenes | 2,712 | 2,247 |
| CDSs (CoDing Sequences) | 97,744 | 71,223 |
| Exons | 526,447 | 444,005 |
| Introns | 467,265 | 381,435 |

## Conclusions

The current genome assembly derived from the Arlee homozygous line and based on long-reads PacBio sequencing provides the most contiguous and complete reference genome generated for rainbow trout to date. It enables a more complete and accurate annotation of protein-coding genes in the genome and can be used for identifying structural genome variations in rainbow trout through comparison with other *de-novo* genome assemblies derived from other rainbow trout genetic lines. We view this assembly as a critical building block towards the generation of a pan-genome reference for rainbow trout.

## Acknowledgements

This project was supported by funds from the USDA Agricultural Research Service in house project number 8082-31000-012 and by the Agriculture and Food Research Initiative Competitive Grant no 2015-07185 from the USDA National Institute of Food and Agriculture. Mention of trade names or commercial products in this publication is solely for the purpose of providing specific information and does not imply recommendation or endorsement by the U.S. Department of Agriculture. USDA is an equal opportunity provider and employer. We thank Roseanna Long and Kristy Shewbridge for their help in the preparation of DNA samples.

## Figure captions

**Figure S1:** The IGHD genes identified in IGHA locus. At the top, a detailed structure of IGHD located at the Cμ-Cδ region. The transmembrane (TM) exons are also indicated. A correspondence between Cδ exons found in Salmo salar (blue boxes) and rainbow trout genome assemblies is shown using dashed lines. At the bottom, the schematic representations of additional IGHD genes (IGHD1 and IGHD2) found in IGHA locus of Arlee genome assembly. The IGHD1 and IGHD2 are upstream of Cμ-Cδ region.

**Figure S2:** Phylogenetic analysis of the Cδ exons annotated at the IGHM-IGHD region of locus IGHA. A phylogenetic tree obtained from the alignment of the Cδ nucleotide sequences from rainbow trout Arlee genome assembly (▴) with those of Salmo salar (▪) and rainbow trout Swanson (●) genome assemblies. The identification of Cδ exons is based on the nomenclature used in Salmo salar and shown in different colors.

**Figure S3**. The amino acid alignment of known rainbow trout secreted IgD sequence and the deduced amino acid sequences from the 10 Cδ exons identified at the IGHM-IGHD region of rainbow trout IGHA locus (Arlee genome assembly). Over the lines the corresponding exons are indicated, following the feature of teleost IgD that it is a hybrid of Cμ1 domain linked to different number of Cδ domains.

**Figure S4:** Phylogenetic analysis of the Cδ exons identified IGHD1 and IGHD2 genes of locus IGHA. The phylogenetic tree obtained from the alignment of the Cδ nucleotide sequences from rainbow trout Arlee genome assembly (▴) with those of *Salmo salar* (▪). The identification of Cδ exons is based on the nomenclature used in *Salmo salar* and shown in different colours.

**Figure S5:** A) The alignment of rainbow trout IgT aminoacid sequences deduced from the functional IGHT genes identified in Arlee genome assembly. Over the lines the corresponding exons are indicated. Consensus sequences, with a threshold >70%, is shown in bold. B) Percentage identity of the amino acid sequence deduced from functional IGHT genes identified in Arlee genome assembly and the three rainbow trout IgT subclasses previously identified (GenBank accession numbers are indicated).

## Supplementary Files

Supplementary File 1: Repeat sequence library generated from the genome sequence.

Supplementary File 2: List of the Hi-C assembly scaffolds.

Supplementary File 3: Graphic presentation of the Hi-C contact map.

Supplementary File 4: Summary statistics of the 32 major scaffolds in the Arlee genome assembly.

Supplementary File 5: Number of scaffolds, contigs and gaps per chromosome in the genome assembly.

Supplementary File 6: SNP haplotypes of the Omy05 double inversion in the 12 WSA doubled haploid lines.

Supplementary File 7: List of small inversions detected between the Swanson and Arlee lines genome assemblies.

